# Rhythmic temperature control in a homeothermic plant: A field study of thermogenic Japanese skunk cabbage, *Symplocarpus renifolius*

**DOI:** 10.1101/2020.04.26.062760

**Authors:** Haruka Tanimoto, Taiga Shibutani, Kota Goto, Kikukatsu Ito

## Abstract

The spadices of skunk cabbage (*Symplocarpus renifolius*) can generate heat and maintain an internal temperature of about 23°C, even when the ambient air temperature drops below freezing. In this study, we continuously measured the spadix temperatures of skunk cabbage together with ambient air temperatures during March 2019 in their natural habitat in northern Honshu on Japan’s main island. Using Fourier transform we showed that the temperature of the thermogenic spadix oscillates over a period of approximately 60 min. Our data suggest that Japanese skunk cabbage possesses a novel timekeeping mechanism that may be associated with a homeothermic control of this plant.

## Introduction

During evolution, plants have developed several mechanisms to cope with adversities such as cold stress. The thermogenesis of skunk cabbage, which belongs to the arum family, is a distinctive example how plants have adapted to the cold environment (Knutson 1974; Seymour & Blaylock 1999; Ito et al. 2003). Japanese skunk cabbage, *Symplocarpus renifolius*, is known to undergo endogenous thermogenesis during flowering (Uemura et al. 1993). Homeothermic thermogenesis of the spadices of skunk cabbage is particularly beneficial to reproductive activities such as pollen tube germination and growth in cold environments (Seymour et al. 2009).

Our previous study suggested that the temperature of thermogenic spadices of skunk cabbage oscillate over a period of about 1 h (Ito et al. 2004) but, to date, no further studies have reported the periodic floral temperature oscillations of skunk cabbage in the field. Because our previous analyses of the periodic temperature fluctuation in thermogenic spadices were conducted using artificially induced and/or shorter-term thermogenic oscillations (Ito et al. 2004), we re-examined these results to determine whether the thermogenic spadix of skunk cabbage inherently exhibits periodic temperature oscillations in their natural habitat in Japan.

## Materials and methods

### Plant materials

The skunk cabbage (*Symplocarpus renifolius*) examined in this study was grown in a field in Iwate Prefecture, Japan (39°14’18.6"N,141°02’51.0"E).

### Temperature measurements and Fourier transformation

Thermogenic spadix and air temperatures were measured at 1-min intervals (Ito et al. 2004; Onda et al. 2008) on May 8, 2019. Fourier analysis was carried out using R software (version 3.6.1).

## Results and discussion

In northern Japan, skunk cabbage blooms in cold environments between late February and early May, when the ground is occasionally covered with snow (Fig. 1a). In the present study, measurements conducted in early March indicated that the maximum and minimum air temperatures were 7.6°C and −2.5°C, respectively (Fig. 1b). In contrast, the spadix temperature appeared to fluctuate, with the average calculated to be 21.7°C; the maximum and minimum temperatures, however, were 25.6°C and 17.7°C, respectively (Fig. 1b). Because pollen germination and the rate of pollen tube growth in skunk cabbage exhibited sharp optima at 23°C (Seymour et al. 2009), the thermogenic spadix studied in this report appeared to be well thermoregulated, both maintaining plant health and enabling fertilization in a cold environment.

**Figure 1.**
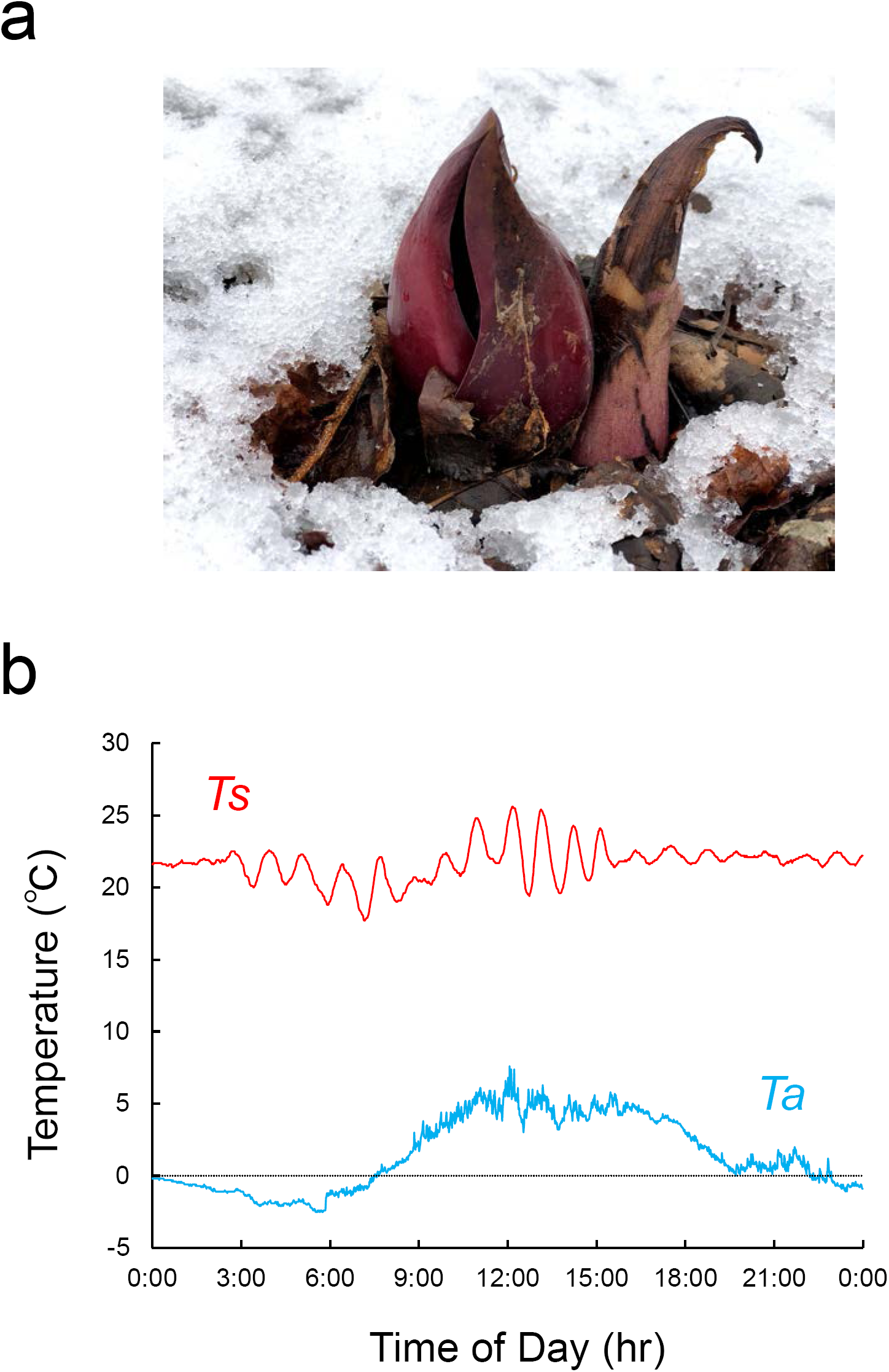
Japanese skunk cabbage, *Symplocarpus renifolius*. (a) Thermogenic stage of skunk cabbage grown outdoors in northern Honshu, on the main island of Japan. The thermogenic spadix is covered with spathe. (b) Temperature changes of the spadix (*Ts*) and ambient air (*Ta*) in the field.

To determine the frequency of the temperature fluctuations in the spadix, we conducted a Fourier analysis using the data obtaining by recording the spadix and ambient air temperatures over one full day, as shown in Fig. 1b. Our results clearly demonstrated that distinct peaks in the frequencies occurred and differed significantly between the spadix (*Ts*) and ambient air (*Ta*) temperatures (Fig. 2a & b). The most dominant peak (peak 7) in the spadix temperature (*Ts*) corresponded to a periodic cycle of 66 min (Table 1). These results are similar to those shown by our previous data, which indicated that artificially induced periodical oscillations of thermogenic spadices of skunk cabbage occurred over an average period of 53.6 ± 4.9 min (Ito et al. 2004). It should be noted here that our previous Fourier analysis involving short-term temperature data demonstrated a periodic cycle of 68.3 min (Ito et al. 2004).

**Table 1.** Frequency of temperature oscillations in the spadix.

| Peak number | Frequency (Hz) | Cycle (min) |
| --- | --- | --- |
| 1 | 1.1566E-05 | 1441 |
| 2 | 1.0409E-04 | 160 |
| 3 | 1.2723E-04 | 131 |
| 4 | 1.5036E-04 | 111 |
| 5 | 1.7349E-04 | 96 |
| 6 | 2.0819E-04 | 80 |
| 7 | 2.5445E-04 | 66 |
| 8 | 3.0072E-04 | 55 |
| 9 | 3.3542E-04 | 50 |
| 10 | 3.8168E-04 | 44 |
| 11 | 4.1638E-04 | 40 |

Note: Peak numbers correspond to those shown in Fig. 2. The major peak (#7) detected in Fig. 2 is shaded.

**Figure 2.**
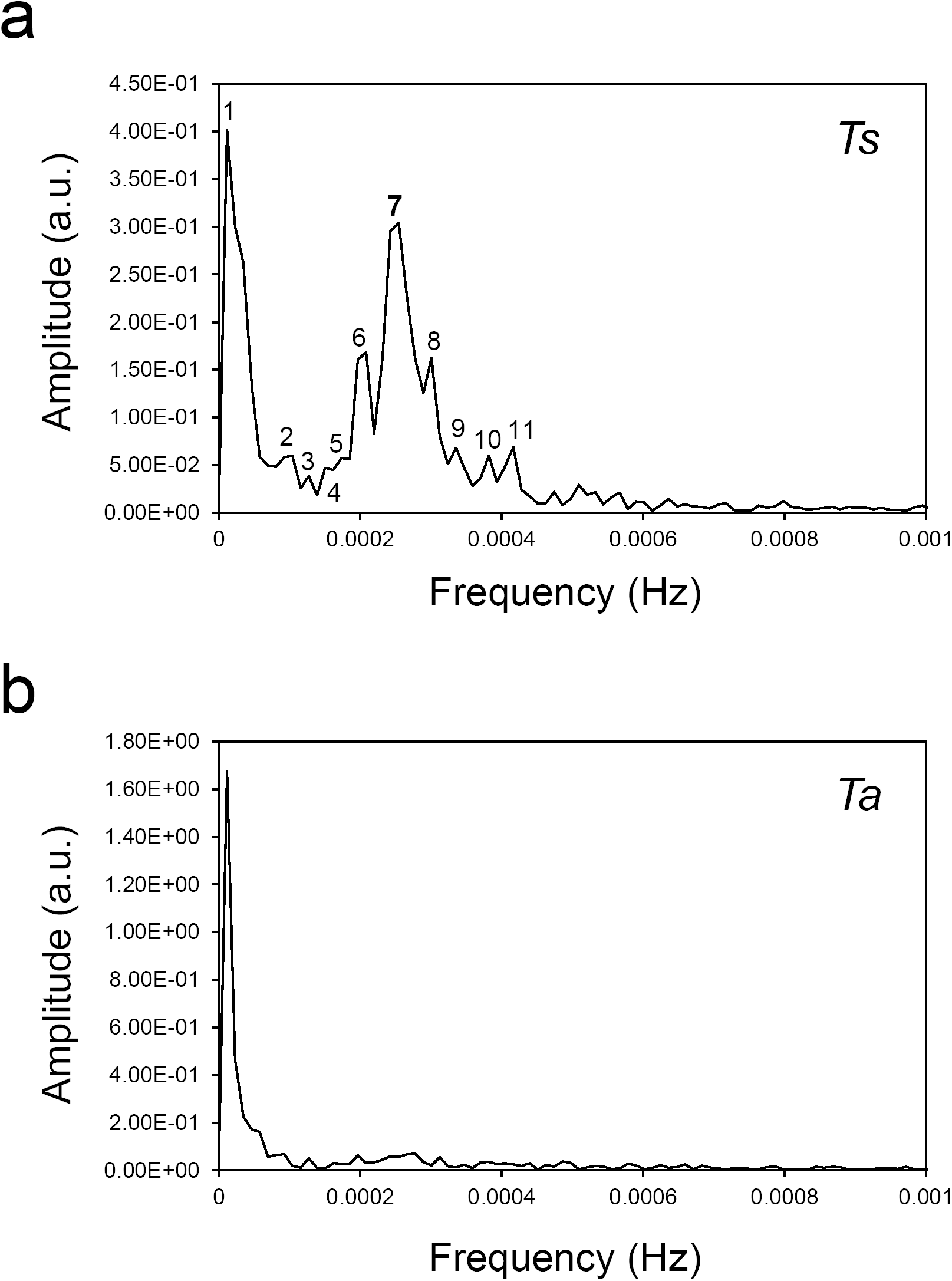
Fourier analysis of temperature data. Analysis involving the temperatures of the spadix (*Ts*) (a) and ambient air (*Ta*) (b). Amplitude is shown as an arbitrary unit (a.u.).

It is plausible that thermogenic spadices of skunk cabbage inherently oscillate their temperatures over a period of approximately 60 min. This would imply that these spadices not only contain a sensitive temperature sensor (Ito et al. 2003; Ito et al. 2004), but also have a biological clock or pacemaker. Presumably, temperature oscillation in skunk cabbage is a novel type of periodic phenomenon that may be included with other well-documented biological rhythms (Goldbeter 1996) (Table 2).

**Table 2.**
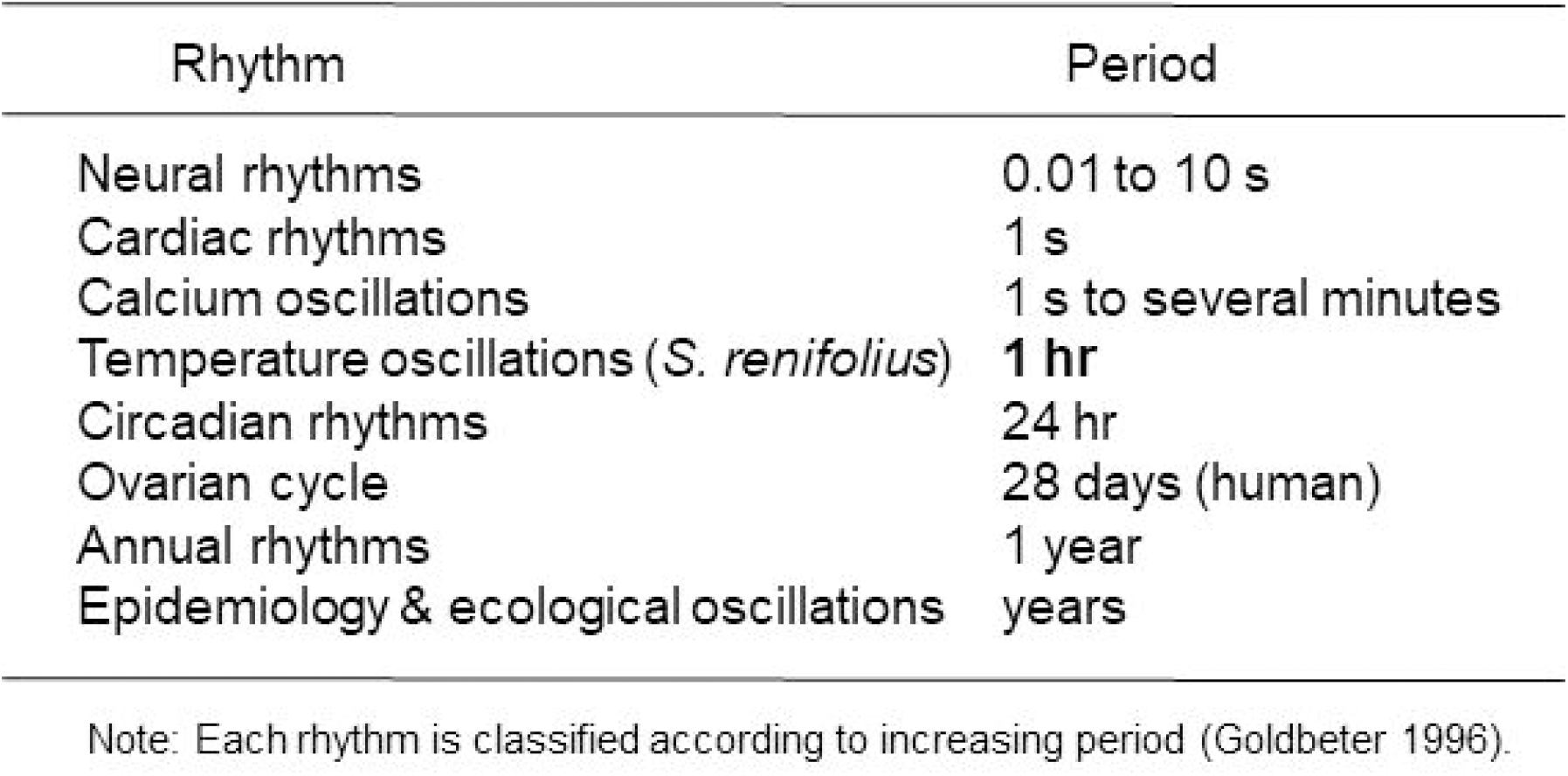
Biological rhythms and periodic cycles of temperature oscillation in skunk cabbage, *Symplocarpus renifo/ius.*

The mechanisms underlying the periodical temperature oscillations determined here remain unclear. It is noteworthy, however, homeothermy of skunk cabbage may be regulated by maintaining thermodynamic equilibrium that involves activation energy that can be adjusted by temperature changes in the thermogenic spadices (Umekawa et al. 2016). It is apparent that changes in temperature influence and shift the point of equilibrium in chemical reactions including those associated with alternative respiration pathways in the spadices of thermogenic skunk cabbage (Moore et al. 2013). A recent report on mammals has indicated that body temperature cycles regulate rhythmic alternative splicing through SR protein phosphorylation (Preußner et al. 2017). It is, therefore, tempting to speculate that the newly identified biological clock or pacemaker in skunk cabbage, which is characterized by a periodic temperature cycle of 60-min, may participate in the thermoregulation of the spadices of skunk cabbage by yet unknown mechanisms. Further investigation is required to identify the detailed mechanisms underpinning periodic temperature regulation in this plant.

## Acknowledgements

The authors are grateful to members of our laboratory for helpful discussions.

## References

Goldbeter A. (1996). Biochemical Oscillations and Cellular Rhythms: The molecular bases of periodic and chaotic behaviour. Cambridge: Cambridge University Press.

Ito K, Ito T, Onda Y, Uemura M. (2004). Temperature-triggered periodical thermogenic oscillations in skunk cabbage (*Symplocarpus foetidus*). Plant and Cell Physiology. 45:257–264.

Ito K, Onda Y, Sato T, Abe Y, Uemura M. (2003). Structural requirements for the perception of ambient temperature signals in homeothermic heat production of skunk cabbage (*Symplocarpus foetidus*). Plant, Cell and Environment. 26:783–788.

Knutson RM. (1974). Heat production and temperature regulation in eastern skunk cabbage. Science. 186:746–747.

Moore AL, Shiba T, Young L, Harada S, Kita K, Ito K. (2013). Unraveling the heater: New insights into the structure of the alternative oxidase. Annu Rev Plant Biol. 64:637–663.

Onda Y, Kato Y, Abe Y, Ito T, Morohashi M, Ito Y, Ichikawa M, Matsukawa K, Kakizaki Y, Koiwa H, Ito K. (2008). Functional coexpression of the mitochondrial alternative oxidase and uncoupling protein underlies thermoregulation in the thermogenic florets of skunk cabbage. Plant Physiol. 146:636–645.

Preußner M, Goldammer G, Neumann A, Haltenhof T, Rautenstrauch P, Müller-McNicoll M, Heyd F. (2017). Body temperature cycles control rhythmic alternative splicing in mammals. Mol Cell. 67:433-446.e434.

Seymour RS, Blaylock AJ. (1999). Switching off the heater: influence of ambient temperature on thermoregulation by eastern skunk cabbage *Symplocarpus foetidus*. J Exp Bot. 50:1525–1532.

Seymour RS, Ito Y, Onda Y, Ito K. (2009). Effects of floral thermogenesis on pollen function in Asian skunk cabbage *Symplocarpus renifolius*. Biol Lett. 5:568–570.

Uemura S, Ohkawara K, Kudo G, Wada N, Higashi S. (1993). Heat-production and cross-pollination of the Asian skunk cabbage *Symplocarpus renifolius* (Araceae). Am J Bot. 80:635–640.

Umekawa Y, Seymour RS, Ito K. (2016). The biochemical basis for thermoregulation in heat-producing flowers. Scientific reports. 6:24830.

